# First detection of a highly invasive freshwater amphipod (*Crangonyx floridanus*) in the United Kingdom

**DOI:** 10.1101/437301

**Authors:** Quentin Mauvisseau, John Davy-Bowker, David Bryson, Graham R. Souch, Alfred Burian, Michael Sweet

**Affiliations:** Aquatic Research Facility, Environmental Sustainability Research Centre, University of Derby, Derby, DE22 1GB, UK; Surescreen Scientifics Ltd, Morley Retreat, Church Lane, Morley, DE7 6DE, UK; Freshwater Biological Association, River Laboratory, East Stoke, Wareham, Dorset, BH20 6BB, UK; Natural History Museum, Cromwell Road, London, SW7 5BD, UK; Forensic Anthropology and Photography, Department of Biomedical and Forensic Science, School of Human Sciences, College of Life and Natural Sciences, University of Derby, DE22 1GB, UK

**Keywords:** *Crangonyx floridanus*, first record, United Kingdom

## Abstract

The freshwater gammarid, *Crangonyx floridanus*, originates from North America but has invaded and subsequently spread rapidly throughout Japan. We provide here the first genetic and microscopic evidence that *C. floridanus* has now also reached the United Kingdom. We found this species in two locations separated by more than 200 km (Lake Windermere in the North of the UK and Smestow Brook, West Midlands). The current distribution of *C. floridanus* is currently unknown, however both sites are well connected to other river and channel systems therefore the chance of further spread is high. Genetic analyses of *C. floridanus* indicate that British inland waters are colonised by the same linage, which also has invaded Japan. We recommend further work to assess the distribution of this species and its impact on the local fauna and flora.

## Introduction

In fresh and brackish waters, amphipods belong to the invertebrate taxa with the highest invasion potential (van der Velde et al., 2009). Non-native amphipods species frequently show high growth rates and may reach carrying capacities substantially above those of local populations (Kotta et al., 2013). This rapid proliferation of invaders leads not only to rapid dispersal but is also frequently linked to dramatic effects on natural communities (Grabowski, Konopacka, Jazdzewski, & Janowska, 2006; Tojo, Tanaka, Kuranishi, & Kanada, 2010; van der Velde et al., 2009). Local populations of gammarid species may be drastically reduced or even replaced completely by the invading species (Grabowski et al., 2006; van der Velde et al., 2009). Furthermore, gammarid amphipods have a varied diet including detritus, plant particles and small invertebrates (Väinölä et al., 2008), coupled with their high grazing rates this (at least in part) explains their impacts when introduced to local ecosystems.

The United Kingdom has already been invaded by at least one *Crangonyx* species, *Crangonyx pseudogracilis* (Bousfield 1958) (also known as *Eucrangonyx gracilis* or *Crangonyx gracilis*). *C. pseudogracilis* originates from North America (Slothouber Galbreath, Smith, Becnel, Butlin, & Dunn, 2010; Zhang & Holsinger, 2003), and was first documented in England in 1936 (Crawford, 1937; Dunn, 2013). Since then, *C. pseudogracilis* has rapidly extended its range. In 2012, it was documented throughout the UK and had colonised various habitats including ponds, lakes, rivers and canal systems (Dunn, 2013; Slothouber Galbreath et al., 2010). However, very little information is available on the species invasion history and its impact on the structure and functionality of local communities (Slothouber Galbreath et al., 2010).

A closely related species *Crangonyx floridanus* (Bousfield 1963) is also originally from the same region of North East America. Similar to *C. pseudogracilis*, *C. floridanus* has a high invasion potential and since 1989 has colonised large regions of the main island of Japan (Nagakubo et al., 2011; Tojo et al., 2010). Its rapid dispersal and high population growth (Nagakubo et al., 2011; Tojo et al., 2010) have highlighted the urgent requirement to contain the spread of *C. floridanus* and protect local species (Nagakubo et al., 2011). Both species are almost identical with field and microscope-based differentiation between the two often resulting in misidentification (Nagakubo et al., 2011). However, to date, there are no records of *C. floridanus* present within Europe (Dunn, 2013). The only visual distinction between these two morphologically similar species is the presence of ventral spines on the outer ramus of the uropod 2 in males *C. pseudogracilis* (Fig. 1), which are absent in males *C. floridanus* (Zhang & Holsinger, 2003). Because of this, a reliable species identification is time consuming and requires substantial taxonomic experience. Differentiation of both species is also possible using comparison of cytochrome C oxidase subunit I (COI) sequences (Nagakubo et al., 2011), which is commonly used in studies of biological invasion of amphipods (Slothouber Galbreath et al., 2010) and in environmental DNA (eDNA) based surveys (Blackman et al., 2017).

**Figure 1.**
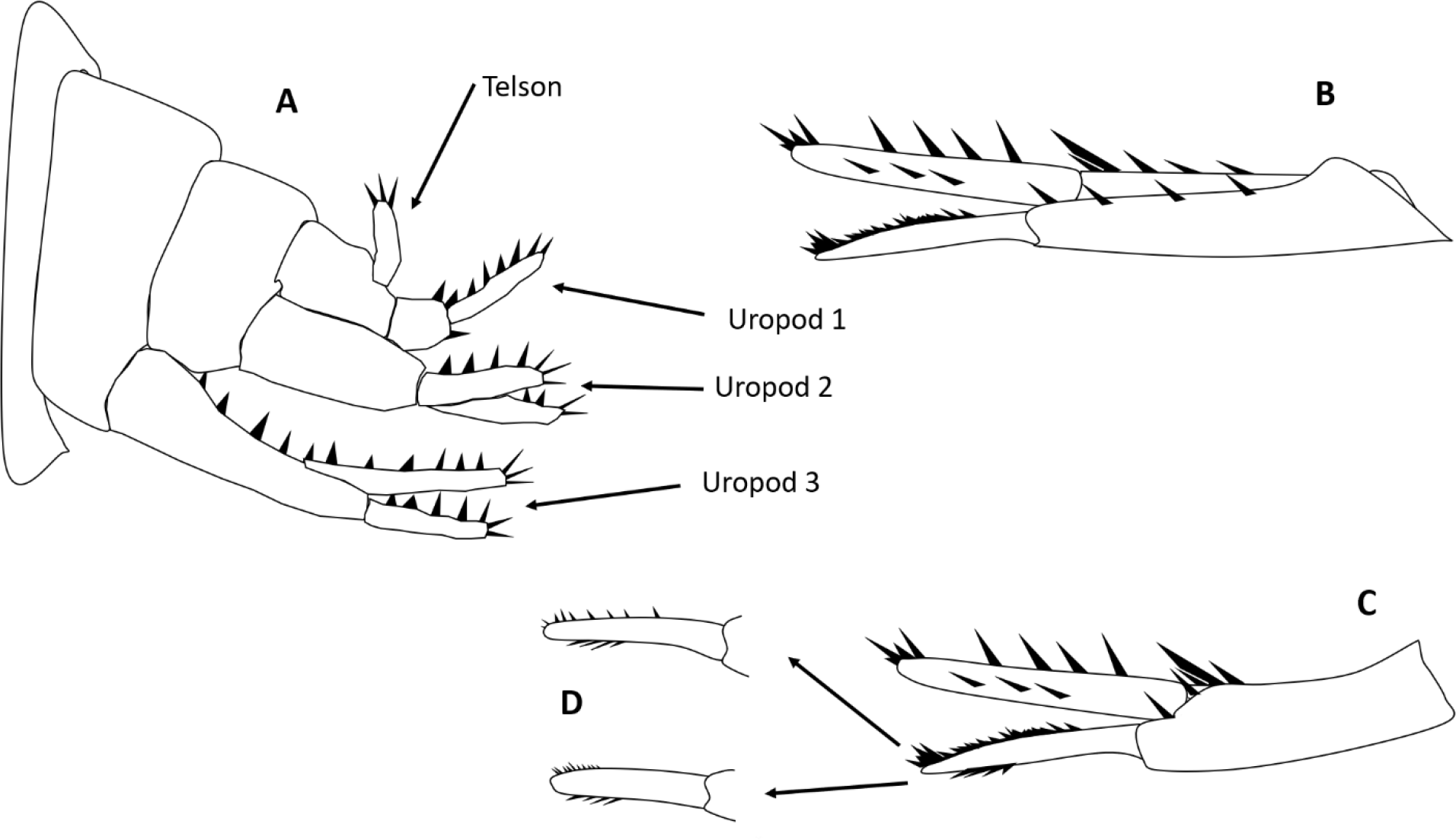
Drawings of the lateral view of posterior abdomen of a *Crangonyx pseudogracilis* specimen (A), uropod 2 of a male *Crangonyx floridanus* specimen (B), uropod 2 of a male *Crangonyx pseudogracilis* specimen (C), with a focus of the outer ramus with ventral spines (D), modified from (Dobson, 2012; Zhang & Holsinger, 2003).

In this paper, we report the first record of *C. floridanus* in Europe (Fig. 2. A). We sampled for the *Crangonyx* species on the 11^th^ September 2017 and again on the 28^th^ July 2018 in Lake Windermere (Latitude: 54.3523 N; Longitude: −2.93880 W) and on 26^th^ September 2018 in the Smestow Brook, West Midlands (Latitude: 52.5727 N; Longitude: −2.2196 W). We present both microscopic and genetic evidence for the presence of *C. floridanus* in these habitats.

**Figure 2.**
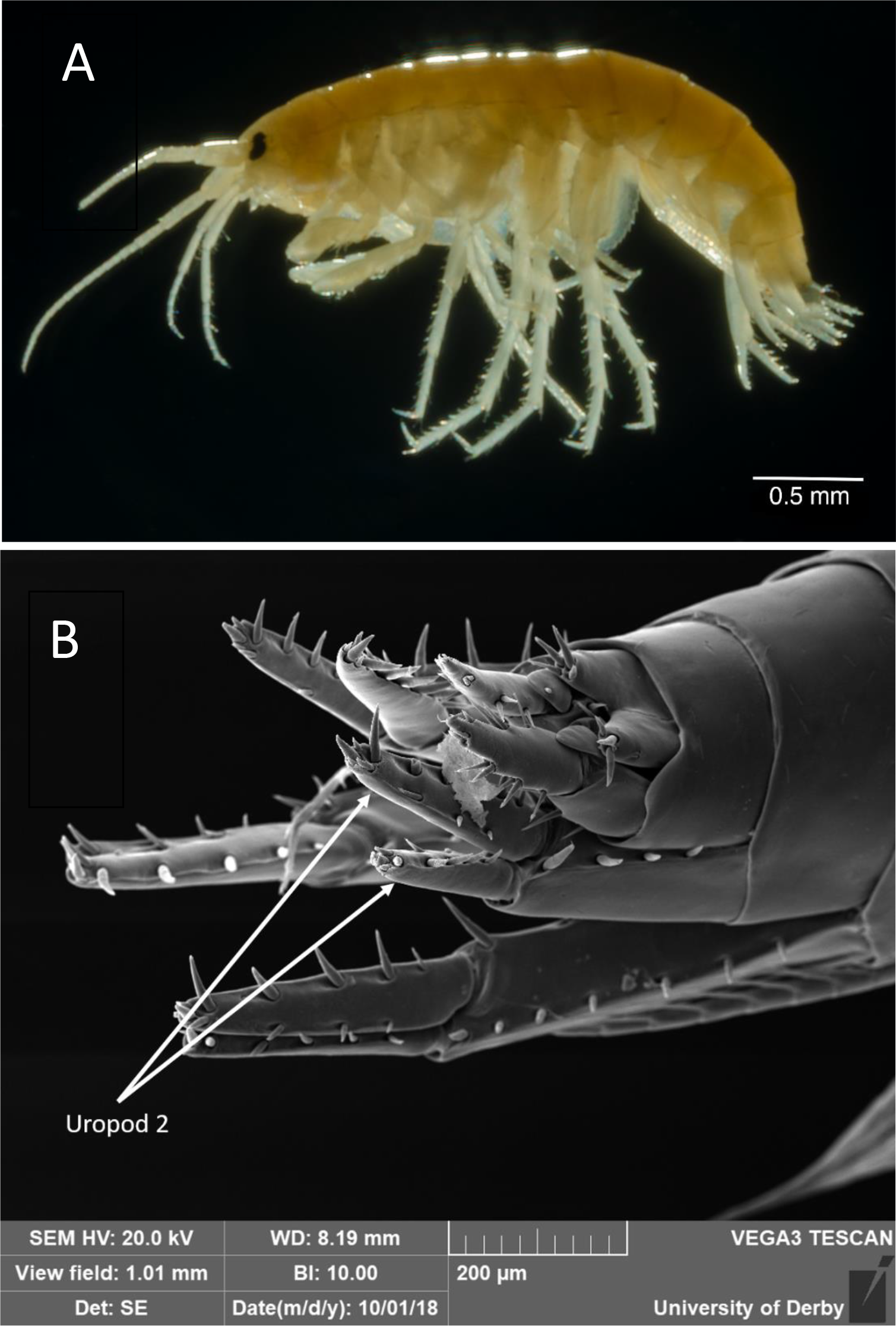
(A) Photomacrography of a *Crangonyx floridanus* specimen found in Windermere Lake, UK, in July 2018. (B) Scanning electron micrograph of the posterior abdomen of a male *Crangonyx floridanus* individual found in Windermere Lake, UK, in July 2018.

## Materials and methods

Lake Windermere is the largest natural lake in England, with a maximal length of 18.08 km, a maximal width of 1.49 km, a maximal depth of 66 m and a surface area of 14.73 km^2^. Lake Windermere is protected as part of the Lake District National Park and the site is actively used for recreation and tourism. Moreover, a station of the British Freshwater Biological Association (FBA) is located close to Lake Windermere and provides facilities for research on the ecology and conservation of local species on the western shore of the lake.

On the 11^th^ September 2017 and later, on the 28^th^ July 2018, kick sampling was undertaken as part of a routine monitoring approach. *Crangonyx* specimens were separated from the samples, preserved in absolute ethanol and transported to the University of Derby for genetic and microscopic analyses. In total, one individual was collected in 2017 and 11 more in 2018. On 26^th^ September 2018, two more Crangonyx specimens were found during kick sampling the submerged root structures of riparian vegetation in Smestow Brook, a small river more than 200 km away from Lake Windermere. Samples were again preserved in 100% ethanol.

### Light and Scanning Electron-Microscopy (SEM)

*Crangonyx* specimens (*n* = 11) were placed on the glass of a Nikon Multiphot diascopic mount and lit by darkfield illumination using a tungsten halogen fibre optic light source. Pictures were taken with a Nikon D7000 mounted on the photomacrography stand, set to colour balance of 3200°K, with a 65mm macro lens (https://www.microscopyu.com/museum/multiphot-large-format-photomacrography-microscope).

Samples for SEM were dehydrated in ethanol, placed on aluminium stubs and coated in gold using a splutter coater (Emitech K550X). Samples were then placed in a Vega 3 SEM (TESCAN) (https://www.tescan.com) and imaged using a SE detector at 20kV.

### Genetic analysis

Prior to performing any lab work, all surfaces were cleaned with 10% and 100% ethanol. All equipment was sterilised for 20 min under UV light before use. All DNA extractions were performed in a PCR free room (i.e. free of any PCR product). DNA extraction were performed using the Qiagen DNeasy Blood and Tissue kit following the manufacturer’s instructions. COI fragments were amplified by PCR using the Forward primer LCO1490 5’-GGTCAACAAATCATAAAGATATTGG-3’ and Reverse primer HCO2198 5’-TAAACTTCAGGGTGACCAAAAAATCA-3’ (Folmer, Black, Hoeh, Lutz, & Vrijenhoek, 1994) as in (Nagakubo et al., 2011). PCR were performed in a 25 μL total volume with 12.5 μL of 2x PCRBIO Ultra Mix Red (PCRBIOSYSTEMS), 1 μL of each primer (10μM), 9.5 μL of ddH2O and 1 μL of DNA template.

The PCR programme was as follow: initial 1 min denaturation at 95°C, 35 cycles of denaturation at 95 °C for 1 min, annealing at 40 °C and elongation at 72 °C for 1 min and 30 s. A final elongation step of 7 min at 72 °C was added at the end of the PCR. Negative controls were added during the PCR to ensure the absence of contamination.

PCR products were confirmed by electrophoresis on 2% agarose gel stained with 3 μL of GelRed^™^ Nucleic Acid Gel Stain, Biotium. Product sizes were checked by comparing amplified DNA to 5 μL of PCRBio Ladder IV (PCRBIOSYSTEMS). PCR product of each specimens was sent for sequencing to the Eurofins Genomics facilities in UK. COI sequences from all 14 specimens were submitted to the NCBI database (https://www.ncbi.nlm.nih.gov/). All sequences were aligned with Geneious 10.1.3 Version and blasted in GenBank for identification of each specimens.

## Results and Discussion

Following the genetic analysis, all 14 individuals from Windermere Lake and Smestow Brook were identified as *C. floridanus* (MK036646, MK036647, MK036648, MK036649, MK036650, MK036651, MK036652, MK036653, MK036654, MK036655, MK036656, MK036657, MK036658, MK036659). Alignment of the sequences showed no variation on the COI fragment between the 14 specimens. All sequences showed 100% similarity with previously uploaded *C. floridanus* sequences (LC171550.1, AB513824.1, AB513818.1, AB513817.1, AB513803.1, AB513801.1, AB513800.1). These have all been sampled in Japan (Nagakubo et al., 2011; Tomikawa, Nakano, Sato, Onodera, & Ohtaka, 2016). Equally, the sequences from this study were all less than 82% similar to a previously published sequence of *C. pseudogracilis* (from the UK-AJ968893.1 -Slothouber Galbreath et al., 2010).

Furthermore, scanning electron micrographs (Fig. 2. B) allowed us to use an alternative approach to distinguish between *C. pseudogracilis* and *C. floridanus* (Zhang & Holsinger, 2003) (Fig. 1). Our analysis clearly demonstrates the absence of ventral spines on the outer ramus of the uropod 2 of male specimen (Fig. 1). Microscopic identification as *C. floridanus* was confirmed for five specimens using SEM and for 11 specimens using light microscopy.

While we provide the first record of *C. floridanus* in the UK., it is unclear whether the species was introduced recently. *C. pseudogracilis* and *C. floridanus* are both small organims i.e. less than 5.5 and 9.0 mm in length for adult males and females respectively (Zhang & Holsinger, 2003). Furthermore, the visual identification of the uropod 2 (used for assessing the absence or presence of ventral spines in male specimens) is almost impossible to do on site (in the field). Consequently, it is certainly possible that individuals have been misidentified for a substantial amount of time. Interestingly, the first specimen of *C. floridanus* collected during our study was mis-identified as a *C. pseudogracilis* by a taxonomic expert before the genetic analysis was undertaken. As the specimens found here (from the UK) are showing 100% similarity with the specimens present in Japan, further studies will be necessary to assess if this specific genotype is especially successful in invasions or if the invasion in UK originated from Japanese individuals in the first place.

The specimens were found in Lake Windermere and in a small brook more than 200 km away. These two geographically distinct sightings suggest that the species has already spread over a substantial area within the UK. The development of an eDNA (i.e. environmental DNA) barcoding approach (Mauvisseau et al., 2018) may represent a future option to easily detect and map both invasive species without any risk of further misidentification. Indeed, eDNA metabarcoding has recently been used to successfully document another invasive gammarid species (Blackman et al., 2017). The authors, then assessed historic specimens and highlighted that this ‘new’ invasive species, *Gammarus fossarum* (Koch 1835) has been present since at least 1964. It is therefore, important to re-assess historic archives using a same methodology identified here to explore if the same is true for *C. floridanus*. Regardless of the timing of any given invasion, it is now an important task to map the distribution of both invasive *Crangonyx* species, assess potential interaction effects of their co-occurrence and evaluate the consequence for native fauna and flora.

## Author contributions

Q.M., M.S., designed the experiment and methodology; J.D.B., Q.M, A.B, performed kick sampling surveys; Q.M., performed genetic analyses; D.B., performed microscopic photography; G.R.S., performed Scanning Electron Microscopy photography. The manuscript was written by Q.M., A.B., and M.S. and reviewed by all authors.

